# Sacral agenesis: A pilot whole exome sequencing and copy number study

**DOI:** 10.1101/058578

**Authors:** Robert M. Porsch, Elisa Merello, Patrizia De Marco, Guo Cheng, Laura Rodriguez, Manting So, Pak C. Sham, Paul K. Tam, Valeria Carpa, Stacey S. Cherny, Maria-Mercè Garcia-Barcelo, Desmond D. Campbell

## Abstract

**Background:** Caudal regression syndrome (CRS) or sacral agenesis is a rare congenital disorder characterized by a constellation of congenital caudal anomalies affecting the caudal spine and spinal cord, the hindgut, the urogenital system, and the lower limbs. CRS is a complex condition, attributed to an abnormal development of the caudal mesoderm, likely caused by the effect of interacting genetic and environmental factors. A well-known risk factor is maternal type 1 diabetes.

**Results:** In this pilot study, exome sequencing and copy number variation (CNV) analyses of 4 CRS trios implicate a number of candidate genes, including *MORN1*, *ZNF330*, *CLTCL1* and *PDZD2*. *De novo* mutations were found in *SPTBN5*, *MORN1* and *ZNF330* and inherited predicted damaging mutations in *PDZD2* (homozygous) and *CLTCL1* (compound heterozygous) as well as in CRS-related genes *PTEN* (heterozygous) and *VANGL1* (heterozygous). In addition, a compound heterozygous mutation in *GLTSCR2*, a direct regulator of *PTEN* was identified.

Two CNV deletions, one *de novo* (chr3q13.13) and one homozygous (chr8p23.2), were detected in one of our CRS patients. These deletions overlapped with CNVs previously reported in patients with similar phenotype.

**Conclusion:** Despite the genetic diversity and the complexity of the phenotype, this pilot study identified genetic features common across CRS patients.

## BACKGROUND

Caudal Regression Syndrome (CRS; Caudal Dysgenesis Syndrome, Caudal Dysplasia Sequence, Congenital Sacral Agenesis; OMIM 600145) is a rare (1 in 7,500-100,000 births [1, 2]) congenital disorder characterized by varying degrees of spinal column agenesis. Associated with it are anomalies of central nervous, genito-urinary, cardiac, respiratory and gastro-intestinal systems [3] with anorectal malformations (ARMs) being the most common.

CRS has been attributed to abnormal fetal development of the caudal mesoderm before the fourth week of gestation [4]. During the abnormal gastrulation, prospective notochordal cells, that are wrongly specified in terms of their rostrocaudal positional encoding, are eliminated. Eventually, fewer or even no cells will be available to form the notochord at a given abnormal segmental level. The consequences of such segmental notochordal paucity are manifold and affect the development of the spinal column and spinal cord as well as other organs that rely on the notochord as their inductor. If the prospective notochord is depleted a wide array of segmental vertebral malformations may develop including segmentation defects, indeterminate or block vertebrae, or absence of several vertebrae. Because of lack of neural induction and absence of a floor plate, fewer prospective neuroectodermal cells will be induced to form the neural tube. The resulting malformation essentially depends on the segmental level and the extent of the abnormality along the longitudinal embryonic axis, with subsequent interference on the processes of primary and/or secondary neurulation [5]. However, what triggers such abnormal events is not known.

Caudal spinal abnormalities are the defining characteristics of CRS. Cama et al. [6] and Pang et al. [7] classified the disorder into 5 categories according to the degree of caudal spine involvement: Type I) total sacral agenesis with normal or short transverse pelvic diameter and some lumbar vertebrae possibly missing; Type II) total sacral agenesis without involvement of lumbar vertebrae; Type III) subtotal sacral agenesis or sacral hypodevelopment; Type IV hemisacrum and Type V) coccygeal agenesis.

Maternal type 1 diabetes is a risk factor for CRS, as it is for many other congenital disorders [8]. Maternal type 1 diabetes confers a higher relative risk (252) for CRS than for any congenital diosorder [9]. The exact mechanism by which maternal diabetes affects fetal development in humans remains unclear [10]. While animal studies have shown that embryos exposed to higher levels of glucose develop growth anomalies, hyperglycemia has not been associated with abnormal fetal development in humans [11]. During normal pregnancies insulin sensitivity is reduced at the start of the third trimester in order to provide metabolic fuel for both mother and the developing fetus. However, since insulin is unable to cross the placenta, the fetus starts producing its own insulin in order to metabolize nutrition. It has been suggested that insulin, antibodies to insulin, or some other abnormality of carbohydrate metabolism could affect the development of a genetically susceptible fetus [12, 13].

Evidence for a genetic cause is provided by the existence of familial segregation as well as animal models. While the most severe forms of CRS present sporadically, milder CRS forms can be transmitted within families in a dominant manner with reduced penetrance and phenotypic variability [4]. Currarino syndrome (CS) is characterized by sacral agenesis type IV, presacral mass, and ARM. CS has been associated with mutations in the *MNX1* gene [14–19]. Yet *MNX1* mutations account for only 50% of sporadic and 90% of familial cases [19]. Although private mutations in genes such as *VANGL1* [20], *HOXD13* [21] and *PTEN* [22] have been described in sporadic cases with caudal dysgenesis and/or vertebrae anomalies, no firm genetic association has been established.

A CRS-like phenotype can be induced by administration in animals of retinoic acid (RA), lithium, cadmium, sulphamide, or organic solvents [23, 24]. Several mutated genes including *Cyp26a1*, *Hoxd13* [25], *Wnt-3a* [26], *Acd*, *Ptf1a*, and *Pcsk5* underlie a CRS-like phenotype in mice [10, 27], yet mutations in the human orthologs have never been identified in CRS patients. Interestingly, the reverse is also true; *Mnx1* (formerly *Hxlb9*) mutant mice do not present Currarino syndrome features [28]. These exceptions to human-mouse phenotypic correlation suggest differences in genetic etiology between humans and experimental organisms [10].

In order to search for genetic risk factors for CRS the exomes of five sporadic CRS cases and their respective healthy parents were sequenced. Due to the sporadic nature of the disease we have focused on *de novo* or recessive inherited damaging genetic variants. In addition we also used a SNP chip assay in order to identify rare and *de novo* CNVs.

## METHODS

### Subjects

The records of patients treated between 1995-2010 at the Neurosurgery Department of Giannina (Genoa, Italy) and at the AbaCid-Genética, Grupo HM Hospitales (Madrid, Spain) for congenital anomalies of the spine were reviewed. For all patients, family history, cardiac, respiratory and endocrine data were collected. This included family history for diabetes. Neurological, neurophysiological (Somatosensory evoked potential, SEP), radiological, neuroradiologic (MRI), orthopaedic, physical, urological (urodynamic, cystography) and surgical assessments were performed for each case. For this pilot study, we selected four Italian trios (CR5, CR17, CR41, CR46) as well as one trio from Spain (CURR20). We only selected cases within this period who had sporadic lower spine agenesis. Patients were also affected with additional anomalies of axial skeleton and internal organs. One child had a mother with diabetes type I. The local ethical committees approved the study and written informed consent was obtained from all patients and parents. Subjects are identified by the trio ID suffixed by either A, B or C indicating father, mother or child respectively.

### Bioinformatics Processing

#### Capture, alignment and base-calling

Whole exome sequencing (WES) was performed at the Centre of Genomic Sciences of the University of Hong Kong, Hong Kong. The exomes of all five trios were sequenced using Illumina HiSeq PE100 and captured with TruSeq Exome Enrichment kit (FC-121-1024, Illumina Inc.). The exome sequences were alignment against Human Genome HG-19 using BWA MEM [29]. Duplicated reads were flagged with Picard-tools [30]. The GATK tool set was used to realign indels, perform base recalibration, remove duplicates, perform indel and SNP calling, and for genotype refinement to improve accuracy of genotype calls [31]. Data quality for each variant was scored and a hard threshold used to remove low data quality variants. We used the GATK recommended criteria for this (see supplementary methods). Relatedness of our participants was investigated using PLINK [32]. We then assessed variants for their potential pathogenicity and frequency, retaining for further analysis only variants that were rare. We considered a variant to be rare if its minor allele frequency was ≤1% in each of several public databases (see supplementary methods). We considered a variant to be potentially deleterious according to a score obtained from KGGSeq [33]. KGGSeq’s prediction algorithm makes use of available biological information (the mutation’s effect on the gene, i.e. stop gain or loss, frameshift, splice site, missense), as well as scoring from other publicly available prediction algorithms (PolyPhen-2, SIFT and others). KGGSeq scores were only used as an informative instrument, variants were not removed from the list of possible disease relevant candidates based on KGGSeq score alone.

#### De novo, homozygous and compound heterozygous mutations

##### Single nucleotide variants (SNVs) and small indels

Subsequent analysis of *de novo* and compound heterozygous, as well as homozygous, mutations was performed using KGGSeq [33]. For recessive disease models (homozygous and compound heterozygous) we only considered variants with a minor allele frequency (MAF) of less than 1%. We defined a *de novo* mutation as a first time genetic alteration of a specific locus in a proband. Compound heterozygous mutations were defined as the co-occurrence of two nonsynonymous alleles, one paternal, the other maternal, within a gene. The probability of *de novo* mutations in each gene was estimated using the framework of Samocha et al. [34].

These probabilities were used to guide the assessment of de novo mutations and not to filter variants. Since a similar framework was not available for compound heterozygous mutations we made use of the only large control trio dataset publicly available, the Genome of the Netherlands (GoNL) [35]. The GoNL is a population dataset containing 250 unaffected parents-offspring trios. We estimated the background compound heterozygous mutation rate per gene from the GoNL dataset. We prioritized compound heterozygous mutations found in our CRS cases in genes where the background rate was low. Thus we classed as a candidate risk locus any gene for which a recessive or *de novo* model could be constructed in any of our trios using the set of rare potentially deleterious variants we had identified. Detected *de novo*, compound heterozygous and homozygous mutations were validated via Sanger sequencing of trio DNA (i. e. genotypes were validated in both parents and child).

Analysis of kinship revealed misattributed paternity within one family (CR46). Hence the family CR46 was excluded from all family based analyses (*de novo*, compound heterozygous and homozygous mutation analysis).

##### Copy number variation

We investigated copy number variation (CNV) in the families CR5, CR17 and CR41 with Illumina’s HumanCoreExome-24 beadchip. Quality control of the assayed genotypes was performed using GenomeStudio (Illumina Inc.) using the default settings. CURR20 was also genotyped using CytoScan® HD Array, but failed initially quality control and was therefore excluded from the analysis. CNV calling and *de novo* CNV detection was performed using PennCNV [36]. We identified potentially disease associating CNVs as follows. We retained for further analysis, CNVs which allowed construction of a recessive disease model for any gene in any of our trios. We also retained *de novo* CNVs and rare CNVs. We deemed a CNV to be rare if it did not overlap with any CNV detected in the 1000 Genome Project. *De novo* CNVs were validated by quantitative real-time PCR (ABI Prism 7900 Sequence Detection System; Applied Biosystems) using TaqMan® Copy Number Assay (Catalog #: 4400291). Ensembl’s genome browser was used to determine genes or regulatory elements affected by the CNVs [37]. Initially we attempted to call CNVs from the exome sequencing data on our trios using three programs (EXCAVATOR [38], CoNIFER[39], and CONTRA [40]), but found no consistency between used tools. Similar previous studies have demonstrated limited power for CNV detection from exome sequencing data tools [41].

## RESULTS

After extensive quality control and MAF (MAF≤1%) filtering we retained 229,849 variants of which 92.4% were known in dbSNP137. Hence we only retained variants below the frequency of 1% and those which are novel. Out of these, 7,442 missense, 184 frameshift, 227 nonframe-shift, 150 splicing, 173 stop-gain and 5 stop-loss variants in 3,872 different genes were analyzed in respect to *de novo*, compound heterozygous and homozygous mutations. Of these variants 236 were either heterozygogous or homozygous in all analyzed cases (see supplementary material II). Out of these, 226 were missense and 10 stop-gain variants distributed across 35 different genes. A further 65 of these variants were predicted to be damaging by KGGSeq. We identified two rare mutations in the known CRS related genes *PTEN* (rs202004587, missense, p.A79T) and *VANGL1* (rs74117015, stop-gain, p.Ser338Ter) in CR5C inherited from the mother and father respectively (Table 2). Both were predicted to be damaging.

**Table 1.** Clinical characteristics of the patients included in this study

| Subjects | Sacral agenesis<sup>1</sup><br>and vertebral<br>malformations | Ribs/Limbs<br>anomalies | Genitourinary | Neural tube | ARM | Cardiac | Other | Maternal<br>phenotype |
| --- | --- | --- | --- | --- | --- | --- | --- | --- |
| CR5/M | CRS<sup>1</sup>-Type II<br>Hemivertebra T7-T8 | Additional 13<sup>th</sup> right rib |  |  |  |  | CPT (II) carrier | CPT (II) deficiency |
| CR17/F | CRS Type II |  | Hydronephrosis<br>Hydroureter<br>Bladder-exstrophy | Lipoma<br>Low-lying conus medullaris |  |  | Omphalocele<br>Twisted teeth | Diabetes type I |
| CR41/F | CRS Type I<br>Deformed T7-T8-T9 | Fusion of 5<sup>th</sup> and 6<sup>th</sup><br>left ribs<br>Additional 13<sup>th</sup> left rib<br>Club feet | Incontinence | Lipoma<br>Blunt ending conus medullaris<br>(T11) |  | Pulmonary<br>vein atresia<br>Inter-ventricular<br>septal defect | Congenital hip<br>dislocation.<br>Motor delay. |  |
| CR46/M | CRS Type II<br>Lumbar kyphosis | Club feet |  |  | Anal<br>Stenosis | Intra-ventricular<br>septal defect | Congenital bilateral<br>hip dislocation.<br>Short neck. | Hydrocephalus |
| CURR20/<br>F | CRS Type V |  |  |  | Anal<br>Stenosis |  | Teratoma |  |

<sup>1</sup>According to Cama et al. [6] and Pang et al. [7] classification of sacral agenesis;. M: male; F: female; T= thoracic vertebra; S: sacral vertebra; ARM: anorectal malformations; CPT: Carnitine palmitoyl transferase

**Table 2.**
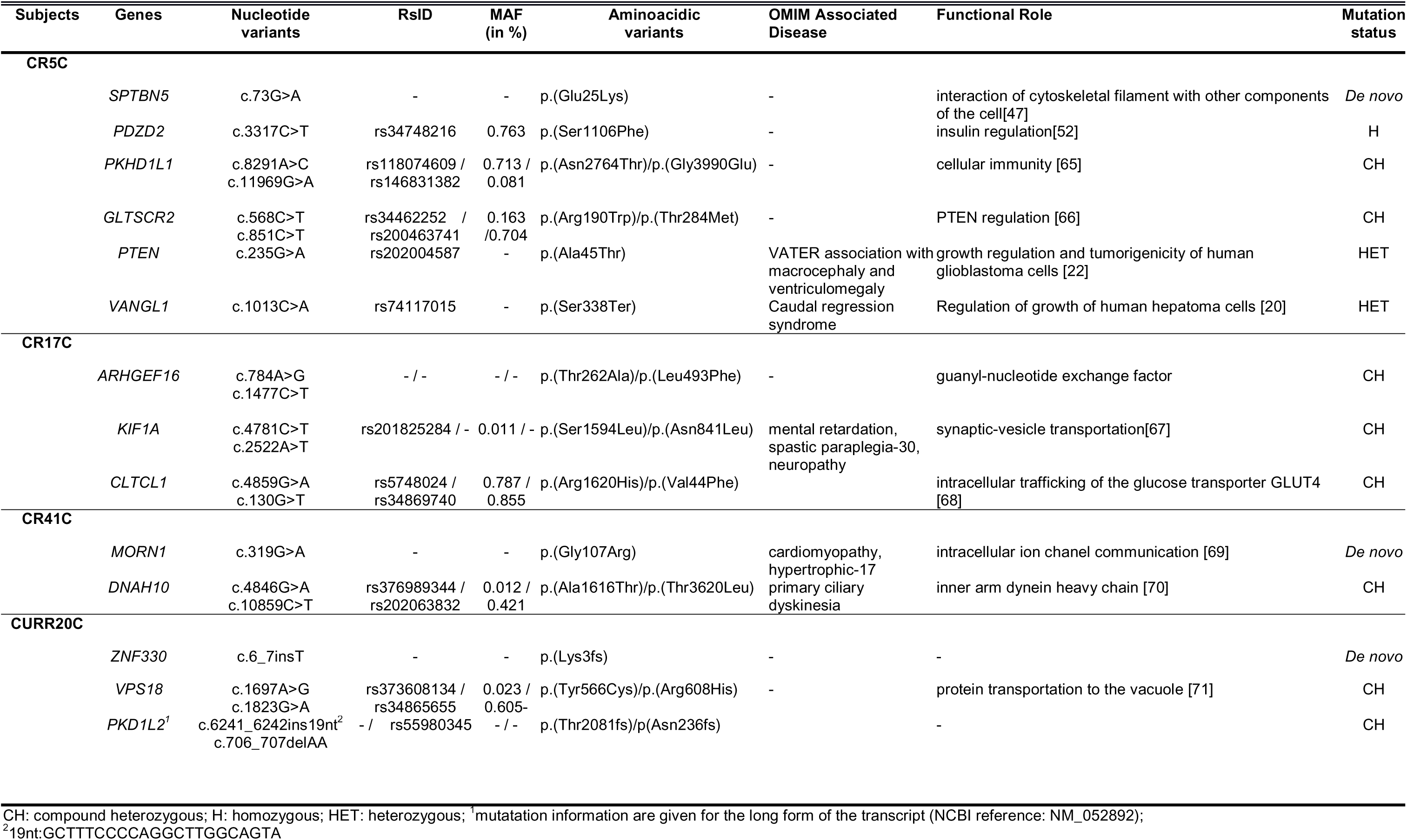
*De novo*, compound heterozygous and homozygous variants.

### De novo variants

In total we identified three *de novo* mutations, two missense and one frameshift mutations in three different genes: *MORN1* (p.Gly107Arg), *SPTBN5* (p.Glu25Lys), and *ZNF330* (p.Lys3fs) in patients CR41C, CR5C, and CURR20C respectively (Table 2). *MORN1* encodes MORN (membrane occupation and recognition nexus) repeats [42]. The exact function of this gene is not known, however, in *Toxoplasma gondii* it is known to be involved in nuclear cell division [43]. Furthermore MORN repeats are known to be part of a number of genes, including junctophilins [44] which are involved in cardiomyopathy [45]. Notably, *MORN1* was reported to be produced by insulin producing cells (IPCs) derived from pancreatic stem cells [46]. The estimated probability for a *de novo* mutation to occur in *MORN1* is 0.8%, but 59% of all analyzed genes have a lower probability [34]. Pathogenicity analysis by KGGSeq suggests that the *de novo* mutation is damaging.

*SPTBN5*(OMIM: 605916) is a beta-spectrin encoding protein. It plays an important role in linking proteins, lipids, and cytosolic factors of the cell membrane to the cytoskeletal filament systems of the cell [47]. *SPTBN5* is expressed in the cerebellum, pancreas, kidney, and bladder, as well as in a number of other systems. The gene has not been associated with any disease or disorder. The estimated gene-specific probability of *de novo* mutation is 1.8% and 99% of genes have a lower probability of having a *de novo* mutation making this this gene less likely to be causally related. Further, KGGSeq’s pathogenic prediction algorithm suggests that this variant is benign.

*ZNF330*(OMIM: 609550) is a zinc finger protein with no known disease association and is mainly present in the nucleus during interphase as well as at the centromeres during mitosis [48]. Interestingly, this gene is differentially expressed in pancreatic Islets of Langerhans and in peripheral blood mononuclear cells [49]. The estimated gene-based *de novo* mutation probability is 0.6%, relatively low but still within the 28th percentile of all genes. KGGSeq was unable to predict a pathogenic score for this variant.

Thus, given the pathogenic nature of the two *de novo* variants and their expression pattern in pancreatic cells, *MORN1* and *ZNF330* are candidate CRS genes.

We detected one *de novo* CNV deletion on 3q13.13 in CR5C (Table 3). The deletion does not seem to encompass any gene or functional element, yet it overlaps with CNVs previously reported in patients with a similar phenotype. In particular, a documented *de novo* deletion in a Japanese patient with OEIS (omphalocele, exstrophy of the cloaca, imperforate anus, spinal defects) complex who also had a sacrum malformation (DECIPHER: 971) overlaps with the *de novo* CNV identified in CR5C.

**Table 3.** *De novo* and homozygous CNVs

| **Patients** | **Chromosome** | **Start Position** | **End Position** | **Length** | **Type** | **Genes or regulatory elements** | **Patients with related symptoms listed in DECIPHER (type of CNV, patient phenotype)** |
| --- | --- | --- | --- | --- | --- | --- | --- |
| **CR5C** |  |  |  |  |  |  |  |
|  | 3q13.13 | 109489534 | 109510473 | 20939 | *De novo*/deletion | - | 971, deletion; abnormality of the sacrum, Abnormality of the small intestine, Anal atresia, Cloacal exstrophy, Omphalocele, Spina bifida occulta. |
|  | 8p23.2 | 5599399 | 5605087 | 5688 | homozygous/deletion | - | 271204, duplication; Abnormality of the sacrum, Central hypotonia, Deeply set eye, Hypermetropia, Long thorax, Narrow mouth, Nasogastric tube feeding in infancy, Seizures, Strabismus. |

### Homozygous and Compound Heterozygous Mutations

In total we identified 8 compound missense heterozygous and one homozygous missense mutations (*PDZD2)* which passed the described filtering criteria (detailed in Table 2). Strikingly, mutations in genes related to diabetes were detected in two patients. None of the affected genes were recurrent. The two genes associated with diabetes were *PDZD2* and *CLTCL1* and were found mutated in CR5C and CR17C respectively.

*PDZD2* (p.Ser1106Phe) has been shown to be an important promoter of fetal pancreatic progenitor cell proliferation [50, 51]. Ma et al. [52] showed that expression of *PDZD2* is specific to pancreatic beta cells. Furthermore, higher concentrations of secreted *PDZD2* in rat insulinoma cell lines were correlated with higher rates of cell proliferation and inhibited transcription of *INS*, an insulin promoter.

*CLTCL1 (*p.Arg1620His, p.Val44Phe) is involved in the intracellular trafficking of glucose transporter *GLUT4*. Intracellular trafficking of the glucose transporter GLUT4 from storage compartments to the plasma membrane is triggered in muscle and fat during the body's response to insulin [53].

A compound heterozygous mutation in *GLTSCR2* (Glioma Tumor Suppressor Candidate Region Gene 2) was identified in patient CR5C (p.Arg190Trp, p.Thr284Met). *GLTSCR2*, is expressed at high levels in pancreas and heart, is a tumor suppressor gene and a direct regulator of *PTEN*. Mutations of *PTEN* have been previously identified in a patient affected with *VACTERL* (Vertebral anomalies, Anal atresia, Cardiac defects, Tracheoesophageal fistula and/or Esophageal atresia, Renal & Radial anomalies and Limb defects) [22] which has commonalities with CRS [27]. This is especially interesting considering that patient CR5C is also harbouring a rare inherited mutation in *PTEN* itself.

A compound heterozygous mutation (p.Ala1616Thr, p.Thr3620Leu) in *DNAH10,* an inner arm dynein heavy chain, was identified in CR41C. Dynein proteins are implicated in many disorders such as motor neuropathies, cortical development diseases, as well as congenital malformations such as heterotaxia and situs inversus. Moreover, cytoplasmic Dyneins have been reported to interact with Kinesin (KIF1A, mutated in patient CR17C) for interkinetic nuclear migration in neural stem cells [54].

We detected a homozygous CNV deletion encompassing part of chromosome 8p23.2 in patient CR5C (Table 3). The CNV does not overlap with known genes but is contained within a duplication found in a patient with abnormal sacrum (DECIPHER: 271204). This documented patient, while also harboring another deletion (7q34-7q36.3), displayed a great variety of phenotypes including central hypertonia, hypermetropia, long thorax, narrow mouth, seizures, strabismus, and deep set eyes. Additional detected rare CNVs overlapped with a number of other genes, however, none were known to be related to CRS (see supplementary material I).

## DISCUSSSION

Within this pilot study we have identified a number of novel risk loci potentially connected to CRS. We have also found a number of mutations for already known genetic risk factors [10]. Here we highlight these preliminary findings and discuss their relevance for future studies.

Foremost, all four patients were affected by a homozygous, compound heterozygous mutations or *de novo* variant in a diabetes-relevant (*CLTCL1* and *PDZD2)* or pancreatic expressed (*MORN1* and *ZNF330)* gene. While these results are not definitive it is in line with the increased CRS risk for children born to diabetic mothers. In addition, one *de novo* (chr3q13.13) and one homozygous CNV (chr8p23.2) overlap with CNVs reported in patients with similar phenotype. Identification of overlapping CNVs in patients with similar phenotype is the central aim of DECIPHER [55]. Since many patients with rare diseases harbor novel or extremely rare variants, it is crucial to accumulate evidence across patients in order to foster understanding of the disease. Furthermore, we identified a heterozygous mutation in *GLTSCR2*, a direct regulator of *PTEN*. *PTEN* has been previously associated with CRS-like phenotypes (VACTER) [10]. Interestingly, the same subject (CR5C) has also an inherited mutation within *PTEN* and the CRS-related gene *VANGL1*. These results further strengthen the role of *PTEN* and *VANGL1* in CRS. Likewise, the identification of compound heterozygous mutations in DNAH10 and KIF1A could suggest an involvement of ciliary proteins.

There are, a number of limitations to our study. Our sample size is small, but to be expected given the disease prevalence and the costs of the genetic assays used. We were not able to identify recurrent affected genes across different patients. We did not look for mutations assuming other than recessive inheritance because the yield of true to false positives would be poor. Many types of genetic variation were not assayed, for instance we only assayed exomic SNVs. Lastly, models involving environment, such as gene-environment interactions within the utero, could not be investigated.

The diversity of identified potential disease mechanisms matches that of previous studies [10, 56–58] and also reflects the phenotypic diversity associated with CRS [56]. We previously showed that if a disorder is genetically complex, one should expect large genetic heterogeneity across patients [59]. Thus the number of candidate genes identified is not surprising and is similar to that reported for other complex rare genetic disorders [60]. While the presence of several possible disease mechanisms does not necessary suggest a multigenic model, the large number of candidate genes identified within this study, as well as those reported by others [23, 27, 61–64] suggests that CRS might be caused by a multitude of private genetic risk factors. This makes identification of a common underlying genetic architecture challenging. Furthermore, differences in the genetic etiology between humans and experimental organisms makes it difficult to investigate the exact causal mechanism. Many aspects of the disease are still poorly characterized, for example, disease prevalence. While some studies have estimated that 1 in 7,700 children might be affected [1], others suggest it might be as rare as 1 in 100,000 births [2]. This further complicates estimation of the number of disease causing mechanisms [59].

### Conclusion

Despite the complexity of the phenotype, we were able to identify common genetic characteristics across patients, potentially causally related to the known risk factors and supposed disease etiology. Our data, although limited to a small group of patients, support a multigenic model for CRS. Future studies should consider larger accumulated samples across multiple centers in order to identify common genetic characteristics via whole genome or whole exome sequencing.

## List of abbreviations

CRS, Caudal Regression; CNV, Copy Number Variation; ARM, Anorectal Malformation; SEP, Somatosensory Evoked Potential; WES, Whole Exome Sequencing; SNP, Single Nucleotide Polymorphism; SNV, Single Nucleotide Variation; MAF, Minor Allele Frequency; IPC, Insulin Producing Cells; Hh, Hedgehog protein; QC, Quality Control

## Availability of data and materials

The corresponding sequencing data, on which this study is based on, can be accessed through the European Genome-phenome Archive (EGA).

## Author’s contributions

RMP, DDC, SSC, GC, MS and MMGB analyzed the data and drafted the paper. VC, EM, PDM and LR played a major role in collecting the samples and phenotyping the patients. MS performed most of the laboratory work. PCS and PKT reviewed the study proposal, provided feedback on the study progress and mansucript. MMGB proposed the study idea.

## Acknowledgements

We thank all participants who made this study possible.

Seed Funding Programme for Basic Research, University of Hong Kong,Project Code:201410159002 to MMGB.

Small Project Funding, University of Hong Kong, Project Code 201209176012 to DC.

EM; PDM, VC would like to acknowledge Ricerca Corrente Ministero della Salute Italia 5X Mille, Aletti- Volpati Trust Onlus and private funding resources. We also thank A.S.B.I. (Associazione Spina Bifida Italia).

## References

1. Unsinn KM, Geley T, Freund MC, Gassner I: US of the spinal cord in newborns: spectrum of normal findings, variants, congenital anomalies, and acquired diseases. Radiographics 2000, 20:923–38.

2. Entezami M: Ultrasound Diagnosis of Fetal Anomalies. Thieme; 2003.

3. Duhamel B: From the Mermaid to Anal Imperforation: The Syndrome of Caudal Regression*. Arch Dis Child 1961, 36:152–155.

4. Barkovich a J, Raghavan N, Chuang S, Peck WW: The wedge-shaped cord terminus: a radiographic sign of caudal regression. AJNR Am J Neuroradiol 1989, 10:1223–31.

5. Tortori-Donati P, Fondelli MP, Rossi a, Raybaud C a, Cama a, Capra V: Segmental spinal dysgenesis: neuroradiologic findings with clinical and embryologic correlation. AJNR Am J Neuroradiol 1999, 20:445–56.

6. Cama A, Palmieri A, Capra V, Piatelli GL, Ravegnani M, Fondelli P: Multidisciplinary management of caudal regression syndrome (26 cases). Eur J Pediatr Surg Off J Austrian Assoc Pediatr Surg. [et al] = Zeitschrift fu r Kinderchirurgie 1996, 6 Suppl 1:44–5.

7. Pang D, Hoffman HJ, Menezes AH: Sacral agenesis and caudal spinal cord malformations. Neurosurgery 1993, 32:755–779.

8. Passarge E, Lenz W: Syndrome of caudal regression in infants of diabetic mothers: observations of further cases. Pediatrics 1966, 37:672–5.

9. Reece EA, Hobbins JC: Diabetic embryopathy: pathogenesis, prenatal diagnosis and prevention. Obstet Gynecol Surv 1986, 41:325–35.

10. Semba K: Etiology of Caudal Regression Syndrome. Hum Genet Embryol 2013, 3.

11. Persaud O: Maternal diabetes and the consequences for her offspring. J Devel Disabil 2007, 13:101–133.

12. Boskovic R, Feig DS, Derewlany L, Knie B, Portnoi G, Koren G: Transfer of insulin lispro across the human placenta: In vitro perfusion studies. Diabetes Care 2003, 26:1390–1394.

13. Challier JC, Hauguel S, Desmaizieres V: Effect of insulin on glucose uptake and metabolism in the human placenta. J Clin Endocrinol Metab 1986, 62:803–7.

14. Hagan DM, Ross a J, Strachan T, Lynch S a, Ruiz-Perez V, Wang YM, Scambler P, Custard E, Reardon W, Hassan S, Nixon P, Papapetrou C, Winter RM, Edwards Y, Morrison K, Barrow M, Cordier-Alex MP, Correia P, Galvin-Parton P a, Gaskill S, Gaskin KJ, Garcia-Minaur S, Gereige R, Hayward R, Homfray T: Mutation analysis and embryonic expression of the HLXB9 Currarino syndrome gene. Am J Hum Genet 2000, 66:1504–15.

15. Garcia-Barceló M, So M-T, Lau DK-C, Leon TY-Y, Yuan Z-W, Cai W-S, Lui VC-H, Fu M, Herbrick J-A, Gutter E, Proud V, Li L, Pierre-Louis J, Aleck K, van Heurn E, Belloni E, Scherer SW, Tam PK-H: Population differences in the polyalanine domain and 6 new mutations in HLXB9 in patients with Currarino syndrome. Clin Chem 2006, 52:46–52.

16. Wang Y, Wu Y: A novel HLXB9 mutation in a Chinese family with Currarino syndrome. Eur J Pediatr Surg 2012, 22:243–5.

17. Ross a J, Ruiz-Perez V, Wang Y, Hagan DM, Scherer S, Lynch S a, Lindsay S, Custard E, Belloni E, Wilson DI, Wadey R, Goodman F, Orstavik KH, Monclair T, Robson S, Reardon W, Burn J, Scambler P, Strachan T: A homeobox gene, HLXB9, is the major locus for dominantly inherited sacral agenesis. Nat Genet 1998, 20:358–61.

18. Kim AY, Yoo S-Y, Kim JH, Eo H, Jeon TY: Currarino syndrome: variable imaging features in three siblings with HLXB9 gene mutation. Clin Imaging 2013, 37:398–402.

19. Belloni E, Martucciello G, Verderio D, Ponti E, Seri M, Jasonni V, Torre M, Ferrari M, Tsui LC, Scherer SW: Involvement of the HLXB9 homeobox gene in Currarino syndrome. Am J Hum Genet 2000, 66:312–9.

20. Kibar Z, Torban E, McDearmid JR, Reynolds A, Berghout J, Mathieu M, Kirillova I, De Marco P, Merello E, Hayes JM, Wallingford JB, Drapeau P, Capra V, Gros P: Mutations in VANGL1 associated with neural-tube defects. N Engl J Med 2007, 356:1432–1437.

21. Garcia-Barceló M-M, Wong KK, Lui VC, Yuan Z, So M, Ngan ES, Miao X, Chung PH, Khong P, Tam PK: Identification of a HOXD13 mutation in a VACTERL patient. Am J Med Genet A 2008, 146A:3181–5.

22. Reardon W, Zhou XP, Eng C: A novel germline mutation of the PTEN gene in a patient with macrocephaly, ventricular dilatation, and features of VATER association. J Med Genet 2001, 38:820–823.

23. Padmanabhan R: Retinoic acid-induced caudal regression syndrome in the mouse fetus. Reprod Toxicol 1998, 12:139–151.

24. Rojansky N, Fasouliotis SJ, Ariel I, Nadjari M: Extreme caudal agenesis. Possible drug-related etiology? J Reprod Med 2002, 47:241–5.

25. Young T, Rowland JE, van de Ven C, Bialecka M, Novoa A, Carapuco M, van Nes J, de Graaff W, Duluc I, Freund J-N, Beck F, Mallo M, Deschamps J: Cdx and Hox genes differentially regulate posterior axial growth in mammalian embryos. Dev Cell 2009, 17:516–26.

26. Greco TL, Takada S, Newhouse MM, McMahon J a, McMahon a P, Camper S a: Analysis of the vestigial tail mutation demonstrates that Wnt-3a gene dosage regulates mouse axial development. Genes Dev 1996, 10:313–324.

27. Szumska D, Pieles G, Essalmani R, Bilski M, Mesnard D, Kaur K, Franklyn A, El Omari K, Jefferis J, Bentham J, Taylor JM, Schneider JE, Arnold SJ, Johnson P, Tymowska-Lalanne Z, Stammers D, Clarke K, Neubauer S, Morris A, Brown SD, Shaw-Smith C, Cama A, Capra V, Ragoussis J, Constam D, Seidah NG, Prat A, Bhattacharya S: VACTERL/caudal regression/Currarino syndrome-like malformations in mice with mutation in the proprotein convertase Pcsk5. Genes Dev 2008, 22:1465–77.

28. Catala M: Genetic control of caudal development. Clin Genet 2002, 61:89–96.

29. Li H: Aligning sequence reads, clone sequences and assembly contigs with BWA-MEM. 2013, 0:3.

30. Picard [http://broadinstitute.github.io/picard/]

31. DePristo MA, Banks E, Poplin R, Garimella K V, Maguire JR, Hartl C, Philippakis AA, del Angel G, Rivas MA, Hanna M, McKenna A, Fennell TJ, Kernytsky AM, Sivachenko AY, Cibulskis K, Gabriel SB, Altshuler D, Daly MJ: A framework for variation discovery and genotyping using next-generation DNA sequencing data. Nat Genet 2011, 43:491–8.

32. Purcell S, Neale B, Todd-Brown K, Thomas L, Ferreira MAR, Bender D, Maller J, Sklar P, de Bakker PIW, Daly MJ, Sham PC: PLINK: a tool set for whole-genome association and population-based linkage analyses. Am J Hum Genet 2007, 81:559–75.

33. Li M-X, Gui H-S, Kwan JSH, Bao S-Y, Sham PC: A comprehensive framework for prioritizing variants in exome sequencing studies of Mendelian diseases. Nucleic Acids Res 2012, 40:e53.

34. Samocha KE, Robinson EB, Sanders SJ, Stevens C, Sabo A, McGrath LM, Kosmicki J a, Rehnström K, Mallick S, Kirby A, Wall DP, MacArthur DG, Gabriel SB, DePristo M, Purcell SM, Palotie A, Boerwinkle E, Buxbaum JD, Cook EH, Gibbs R a, Schellenberg GD, Sutcliffe JS, Devlin B, Roeder K, Neale BM, Daly MJ: A framework for the interpretation of de novo mutation in human disease. Nat Genet 2014(August).

35. The Genome of the Netherlands Consortium: Whole-genome sequence variation, population structure and demographic history of the Dutch population. Nat Genet 2014.

36. Wang K, Li M, Hadley D, Liu R, Glessner J, Grant SF a, Hakonarson H, Bucan M: PennCNV: An integrated hidden Markov model designed for high-resolution copy number variation detection in whole-genome SNP genotyping data. Genome Res 2007, 17:1665–1674.

37. Cunningham F, Amode MR, Barrell D, Beal K, Billis K, Brent S, Carvalho-Silva D, Clapham P, Coates G, Fitzgerald S, Gil L, Girón CG, Gordon L, Hourlier T, Hunt SE, Janacek SH, Johnson N, Juettemann T, Kähäri AK, Keenan S, Martin FJ, Maurel T, McLaren W, Murphy DN, Nag R, Overduin B, Parker A, Patricio M, Perry E, Pignatelli M, et al.: Ensembl 2015. Nucleic Acids Res 2014, 43:D662–669.

38. Magi A, Tattini L, Cifola I, D’Aurizio R, Benelli M, Mangano E, Battaglia C, Bonora E, Kurg A, Seri M, Magini P, Giusti B, Romeo G, Pippucci T, De Bellis G, Abbate R, Gensini GF: EXCAVATOR: detecting copy number variants from whole-exome sequencing data. Genome Biol 2013, 14:R120.

39. Krumm N, Sudmant PH, Ko A, O’Roak BJ, Malig M, Coe BP, Quinlan AR, Nickerson DA, Eichler EE: Copy number variation detection and genotyping from exome sequence data. Genome Res 2012, 22:1525–1532.

40. Li J, Lupat R, Amarasinghe KC, Thompson ER, Doyle MA, Ryland GL, Tothill RW, Halgamuge SK, Campbell IG, Gorringe KL: CONTRA: Copy number analysis for targeted resequencing. Bioinformatics 2012, 28:1307–1313.

41. Tan R, Wang Y, Kleinstein SE, Liu Y, Zhu X, Guo H, Jiang Q, Allen AS, Zhu M: An evaluation of copy number variation detection tools from whole-exome sequencing data. Hum Mutat 2014, 35:899–907.

42. Gerhard DS, Wagner L, Feingold EA, Shenmen CM, Grouse LH, Schuler G, Klein SL, Old S, Rasooly R, Good P, Guyer M, Peck AM, Derge JG, Lipman D, Collins FS, Jang W, Sherry S, Feolo M, Misquitta L, Lee E, Rotmistrovsky K, Greenhut SF, Schaefer CF, Buetow K, Bonner TI, Haussler D, Kent J, Kiekhaus M, Furey T, Brent M, et al.: The status, quality, and expansion of the NIH full-length cDNA project: the Mammalian Gene Collection (MGC). Genome Res 2004, 14:2121–7.

43. Ferguson DJP, Sahoo N, Pinches RA, Bumstead JM, Tomley FM, Gubbels M-J: MORN1 has a conserved role in asexual and sexual development across the apicomplexa. Eukaryot Cell 2008, 7:698–711.

44. Takeshima H, Komazaki S, Nishi M, Iino M, Kangawa K: Junctophilins: a novel family of junctional membrane complex proteins. Mol Cell 2000, 6:11–22.

45. Matsushita Y, Furukawa T, Kasanuki H, Nishibatake M, Kurihara Y, Ikeda A, Kamatani N, Takeshima H, Matsuoka R: Mutation of junctophilin type 2 associated with hypertrophic cardiomyopathy. J Hum Genet 2007, 52:543–8.

46. Ebrahimie M, Esmaeili F, Cheraghi S, Houshmand F, Shabani L, Ebrahimie E: Efficient and simple production of insulin-producing cells from embryonal carcinoma stem cells using mouse neonate pancreas extract, as a natural inducer. PLoS One 2014, 9.

47. Stabach PR, Morrow JS: Identification and characterization of beta V spectrin, a mammalian ortholog of Drosophila beta H spectrin. J Biol Chem 2000, 275:21385–95.

48. Bolívar J, Díaz I, Iglesias C, Valdivia MM: Molecular cloning of a zinc finger autoantigen transiently associated with interphase nucleolus and mitotic centromeres and midbodies. Orthologous proteins with nine CXXC motifs highly conserved from nematodes to humans. J Biol Chem 1999, 274:36456–64.

49. Klijn C, Durinck S, Stawiski EW, Haverty PM, Jiang Z, Liu H, Degenhardt J, Mayba O, Gnad F, Liu J, Pau G, Reeder J, Cao Y, Mukhyala K, Selvaraj SK, Yu M, Zynda GJ, Brauer MJ, Wu TD, Gentleman RC, Manning G, Yauch RL, Bourgon R, Stokoe D, Modrusan Z, Neve RM, de Sauvage FJ, Settleman J, Seshagiri S, Zhang Z: A comprehensive transcriptional portrait of human cancer cell lines. Nat Biotechnol 2014, 33:306–12.

50. Tsang SW, Shao D, Cheah KSE, Okuse K, Leung PS, Yao K-M: Increased basal insulin secretion in Pdzd2-deficient mice. Mol Cell Endocrinol 2010, 315:263–70.

51. Suen PM, Zou C, Zhang YA, Lau TK, Chan J, Yao KM, Leung PS: PDZ-domain containing-2 (PDZD2) is a novel factor that affects the growth and differentiation of human fetal pancreatic progenitor cells. Int J Biochem Cell Biol 2008, 40:789–803.

52. Ma RYM, Tam TSM, Suen APM, Yeung PML, Tsang SW, Chung SK, Thomas MK, Leung PS, Yao K-MM: Secreted PDZD2 exerts concentration-dependent effects on the proliferation of INS-1E cells. Int J Biochem Cell Biol 2006, 38:1015–1022.

53. Vassilopoulos S, Esk C, Hoshino S, Funke BH, Chen C-Y, Plocik AM, Wright WE, Kucherlapati R, Brodsky FM: A role for the CHC22 clathrin heavy-chain isoform in human glucose metabolism. Science 2009, 324:1192–6.

54. Tsai J-W, Lian W-N, Kemal S, Kriegstein AR, Vallee RB: Kinesin 3 and cytoplasmic dynein mediate interkinetic nuclear migration in neural stem cells. Nat Neurosci 2010, 13:1463–1471.

55. Firth H V, Richards SM, Bevan AP, Clayton S, Corpas M, Rajan D, Van Vooren S, Moreau Y, Pettett RM, Carter NP, Lu X, Shaw CA, Patel A, Li J, Cooper ML, Wells WR, Sullivan CM, Sahoo T, Yatsenko SA, Bacino CA, al. et, Shaw-Smith C, Redon R, Rickman L, Rio M, Willatt L, Fiegler H, Firth H, Sanlaville D, Winter R, et al.: DECIPHER: Database of Chromosomal Imbalance and Phenotype in Humans Using Ensembl Resources. Am J Hum Genet 2009, 84:524–33.

56. Singh SK, Singh RD, Sharma A: Caudal regression syndrome--case report and review of literature. Pediatr Surg Int 2005, 21:578–81.

57. Kaciński M, Jaworek M, Skowronek-Bała B: Caudal regression syndrome associated with the white matter lesions and chromosome 18p11.2 deletion. Brain Dev 2007, 29:164–6.

58. Chan BWH, Chan K-S, Koide T, Yeung S-M, Leung MBW, Copp AJ, Loeken MR, Shiroishi T, Shum ASW: Maternal diabetes increases the risk of caudal regression caused by retinoic acid. Diabetes 2002, 51:2811–6.

59. Campbell DD, Porsch RM, Cherny SS, Capra V, Merello E, De Marco P, Sham PC, Garcia-Barceló M-M: Cost effective assay choice for rare disease study designs. Orphanet J Rare Dis 2015, 10:10.

60. Boycott KM, Vanstone MR, Bulman DE, MacKenzie AE: Rare-disease genetics in the era of next-generation sequencing: discovery to translation. Nat Rev Genet 2013, 14:681–691.

61. Adra A, Cordero D, Mejides A, Yasin S, Salman F, O’Sullivan MJ: Caudal regression syndrome: etiopathogenesis, prenatal diagnosis, and perinatal management. Obstet Gynecol Surv 1994, 49:508–16.

62. Vlangos C, O’Connor B, Morley M: Caudal regression in adrenocortical dysplasia (acd) mice is caused by telomere dysfunction with subsequent p53-dependent apoptosis. Dev … 2009, 334:418–428.

63. Knight B: Caudal Regression Syndrome: A Case Report. AANA J 2011, 79:281–282.

64. Evans J a, Vitez M, Czeizel a: Congenital abnormalities associated with limb deficiency defects: a population study based on cases from the Hungarian Congenital Malformation Registry (1975-1984). Am J Med Genet 1994, 49:52–66.

65. Hogan MC, Griffin MD, Rossetti S, Torres VE, Ward CJ, Harris PC: PKHDL1, a homolog of the autosomal recessive polycystic kidney disease gene, encodes a receptor with inducible T lymphocyte expression. Hum Mol Genet 2003, 12:685–698.

66. Yim J-H, Kim Y-J, Ko J-H, Cho Y-E, Kim S-M, Kim J-Y, Lee S, Park J-H: The putative tumor suppressor gene GLTSCR2 induces PTEN-modulated cell death. Cell Death Differ 2007, 14:1872–9.

67. Rivire JB, Ramalingam S, Lavastre V, Shekarabi M, Holbert S, Lafontaine J, Srour M, Merner N, Rochefort D, Hince P, Gaudet R, Mes-Masson AM, Baets J, Houlden H, Brais B, Nicholson G a., Van Esch H, Nafissi S, De Jonghe P, Reilly MM, Timmerman V, Dion P a., Rouleau G a.: KIF1A, an axonal transporter of synaptic vesicles, is mutated in hereditary sensory and autonomic neuropathy type 2. Am J Hum Genet 2011, 89:219–301.

68. Nahorski MS, Al-Gazali L, Hertecant J, Owen DJ, Borner GHH, Chen Y-C, Benn CL, Carvalho OP, Shaikh SS, Phelan A, Robinson MS, Royle SJ, Woods CG: A novel disorder reveals clathrin heavy chain-22 is essential for human pain and touch development. Brain 2015, 138:2147–2160.

69. Landstrom AP, Weisleder N, Batalden KB, Bos JM, Tester DJ, Ommen SR, Wehrens XHT, Claycomb WC, Ko J-K, Hwang M, Pan Z, Ma J, Ackerman MJ: Mutations in JPH2-encoded junctophilin-2 associated with hypertrophic cardiomyopathy in humans. J Mol Cell Cardiol 2007, 42:1026–35.

70. Maiti AK, Mattéi MG, Jorissen M, Volz A, Zeigler A, Bouvagnet P: Identification, tissue specific expression, and chromosomal localisation of several human dynein heavy chain genes. Eur J Hum Genet 2000, 8:923–32.

71. Huizing M, Didier A, Walenta J, Anikster Y, Gahl W a., Krämer H: Molecular cloning and characterization of human VPS18, VPS 11, VPS16, and VPS33. Gene 2001, 264:241–247.

